# CLCFM3: An advanced photogrammetry algorithm for high-precision plant 3D modeling

**DOI:** 10.1101/2024.10.10.617704

**Authors:** Atsushi Hayashi, Nobuo Kochi, Kunihiro Kodama, Sachiko Isobe, Takanari Tanabata

## Abstract

This research aims to develop a novel technique to acquire a large amount of high-density, high-precision 3D point cloud data for plant phenotyping using photogrammetry technology. The complexity of plant structures, characterized by overlapping thin parts such as leaves and stems, makes it difficult to reconstruct accurate 3D point clouds. One challenge in this regard is occlusion, where points in the 3D point cloud cannot be obtained due to overlapping parts, preventing accurate point capture. Another is the generation of erroneous points in non-existent locations due to image-matching errors along object outlines. To overcome these challenges, we propose a 3D point cloud reconstruction method, CLCFM3 (Closed-Loop Coarse-to-Fine method with Multi-Masked Matching). This method repeatedly executes a process that generates point clouds locally to suppress occlusion (multi-matching) and a process that removes noise points using a mask image (masked matching). Furthermore, we propose the Closed-Loop Coarse-to-Fine Method (CLCFM) to improve the accuracy of structure from motion, which is essential for implementing the proposed point cloud reconstruction method. CLCFM solves loop-closure by performing coarse-to-fine camera position estimation. By facilitating the acquisition of high-density, high-precision 3D data on a large number of plant bodies, as is necessary for research activities, this approach is expected to enable comparative analysis of visible phenotypes in the growth process of a wide range of plant species based on 3D information.

## 1. Introduction

The phenotypes of plants and plant organs are important information in plant functional analyses. With the development of molecular biology, it is becoming increasingly necessary to obtain large quantities of highly accurate data. Digital imaging technology is a simple, low-cost method that offers the advantage of nondestructive measurements of samples [1]. However, the images captured by cameras are 2D information, and there are limits to how accurately 2D cameras can obtain 3D shapes and structures, including depth. To precisely measure plant phenotypes during the growth process, technology is needed to accurately digitize shapes and structures in 3D space. The development of measurement technology to support this is progressing [2, 3, 4].

Two main types of measurement technology are used to acquire 3D information about an object. One is the active method, which uses a device that emits light such as laser light to perform measurements, and the other is the passive method, which reconstructs 3D data from photographed images of the target [5]. An example of the latter is photogrammetry SfM/MVS (Structure from Motion/Multi-View Stereo) [6]. Photogrammetry can be implemented using commercially available digital cameras. It offers two following advantages. First, the hardware cost is low. Second the necessary measurements can be taken simply by photographing the target from all directions, so a flexible measurement system can be constructed according to the measurement environment [7]. To obtain phenotypic data for functional analysis, it is necessary to measure different plant bodies of varying sizes and structural complexities, to perform measurements in various locations, and to measure large numbers of individuals. Photogrammetry can solve these issues.

Photogrammetry is a method of reconstructing 3D point cloud data of an object using images taken from various angles. This is realized by combining SfM processing and MVS processing. SfM processing arranges the viewpoint positions (hereafter referred to as camera positions) of images taken in three-dimensional space [8, 9]. MVS processing searches for the position in the image that corresponds to the target point and combines this with the camera position information obtained by SfM to ascertain the position of the point in three-dimensional space using stereo matching processing. This process is repeated to reconstruct a dense 3D point cloud of the object in the image [10]. Given its ability to generate dense 3D point clouds from images, this approach has been proposed for use in phenotyping [11].

Plant structures are characterized by the overlapping of thin, narrow parts such as stems and leaves. Thus, image matching using stereo matching is often unsuccessful because of occlusion. Image matching is also likely to fail because leaf and stem surfaces are similar in color and texture. These failures in image matching can lead to problems in estimating the camera position and can result in incomplete sections of the plant body within the sought 3D point cloud [12] or the occurrence of noise points in places other than the plant body [13]. Creating a plant 3D model with low accuracy is relatively easy, but it remains challenging to construct high-precision 3D models at high throughput for measurement purposes. Therefore, when using photogrammetry to measure plants, it is necessary to solve the problem of failures in image matching that occur in SfM and MVS processing.

To address such issues in SfM processing, various proposals have been made to enhance the accuracy of camera position estimation. These methods involve integrating reference points or GPS devices to provide positional information during capture, such as utilizing images taken with 3D reference points placed around the object to be measured [14, 15] or employing positional information obtained from GPS [16]. On the other hand, as a solution in image processing, the loop closing method has been proposed. This method minimizes estimation errors by linking the initial image captured with the final image when all surrounding images of an object are taken [17]. In the image matching process during camera position estimation, techniques have been proposed to pre-evaluate the entire image’s similarity and the relation between images. This approach aims to mitigate the buildup of matching errors resulting from similar patterns [18, 19, 20].

A major issue with MVS processing is the creation of noise points. Zhang et al. reconstructed 3D models of sweet potato and paprika using SfM/MVS. They found that the 3D reconstruction failure was caused by the inaccuracy of background removal and shadows caused by lighting. Additionally, they noted that the lack of effective solutions for matching at the boundaries of the contours between the background and the object contributes to the inaccuracy of 3D reconstruction for plants [3, 21]. Some inaccuracies are projected as noise points. One reason for the presence of noise points is the occurrence of “flying pixels.” These isolated points emerge at object boundaries where there is a depth change between objects. They are considered noise points as they appear in locations devoid of actual objects. Flying pixels occur more prominently in active methods such as laser scanners, LiDAR, and ToF cameras than in passive methods. Methods for removing them utilize the characteristics of the scanning data employed in active methods. The problem of flying pixels has been confirmed not only in active methods but also in passive methods that use images, such as stereo cameras [22]. The noise points of flying pixels in the passive method result from errors in the stereo matching process around the edges of the measurement target. In particular, the matching results become unstable at the boundary areas where the object and background are mixed in one pixel of the image, and noise points are generated according to these matching results. However, it is difficult to theoretically distinguish whether these points are background or object and to remove them in post-processing. This problem remains unresolved given the fundamental disparity between passive and active methods. Therefore, many methods have been proposed for plant research, with most of them involving the removal of noise points from the 3D point cloud data after reconstruction for post-processing [23]. That is, the noise in the point cloud data is solved by filtering it out in post-processing. For example, one proposed method uses the Euclidean clustering algorithm to segment the plant point cloud data and remove irrelevant background as well as spatially discontinuous points [24].

This research aims to acquire 3D point clouds of individual plants for phenotypic evaluation of plant organ shapes, individual structures, and so on. To this end, we have developed a technology that uses photogrammetry to acquire a large amount of high-density, high-precision 3D point cloud data of plants. Non-photogrammetric methods for reconstructing 3D point clouds using images have also been proposed, with deep learning‒based methods such as NeRF [25] being representative. These methods entail training costs and, when high-resolution images are used as input images, a large amount of flying pixel noise [26]. Moreover, the results have been confirmed to differ from the original shape of the object, which has uneven surfaces on smooth planes and smooth curved surfaces [27]. For this reason, these methods are not suitable for the acquisition of 3D point cloud data, which is our objective. Furthermore, 3D point clouds with a spatial resolution of 1 mm for plants have been reconstructed by utilizing photogrammetry techniques [28], demonstrating the effectiveness of the use of photogrammetry.

This paper proposes a method, CLCFM3, for solving the problems of photogrammetry for plants. CLCFM3 combines two newly proposed methods: MMM and CLCFM. MMM (Multi-Masked Matching) combines the suppression of occlusion by locally generating point clouds (Multi-Matching) using images with adjacent camera positions and Masked Matching, which removes flying pixel noise that tends to occur at the boundary of the target object. Multi-matching generates a 3D point cloud with reduced occlusion defects and then performs a noise point deletion process using images taken from different angles surrounding those utilized in multi-matching. Masked matching effectively reduces flying pixel noise, which has been considered difficult to remove until now. To perform MMM processing, it is necessary to improve the accuracy of the camera position estimation using SfM processing. To achieve this, we proposed CLCFM, which suppresses the accumulation of camera position estimation errors. CLCFM3, which integrates MMM and CLCFM, can reconstruct high-density 3D point clouds with fewer noise points and fewer missing points. We created an analysis pipeline that implements CLCFM3 and applied the point cloud reconstruction process to soybeans. The results showed that it is possible to reconstruct point clouds with fewer missing points by reducing the noise caused by flying pixels. Moreover, the measurement accuracy achieved with this method revealed that the 3D models constructed through it possess sufficient precision for plant measurement. This suggests that the proposed method holds the potential to truly enable the application of 3D models for precise plant measurements.

## 2. Materials and Methods

### 2.1. Plant material and cultivation

A total of four varieties were used: three soybean varieties—GmJMC092 (KUROHIRA), GmJMC112 (FUKUYUTAKA), and HOJAK—were derived from the Genebank project, NARO (https://www.gene.affrc.go.jp), while the fourth variety, MISUZUDAIZU, was derived from the NBRP Lotus japonicus & Glycine max collection (https://nbrp.jp/en/resource/lotus-glycine-en/). The seeds were sown in 1/5000a Wagner pots in 2018 using baked soil, and the plants were grown in a greenhouse at the Kazusa DNA Research Institute. Water tubes were installed in the pots, with irrigation occurring twice a day.

### 2.2. *CLCFM3*: 3D reconstruction algorithm

We propose a 3D reconstruction method called CLCFM3, based on all-round images of a rotating object taken in front of a camera, assuming that each image is accompanied by information on the order in which it was taken. CLCFM3 consists of two processes: MMM and CFCFM, which utilize information on the order in which images were taken. MMM selects adjacent images and performs MVS processing locally to reduce the number of defects caused by occlusion. CLCFM first selects a part of the captured image and performs SfM before iteratively adding portions of the captured image to the SfM results for further processing. An important point of the proposed method is that it selects images using order-of-capture information so as to surround the measurement target, thus minimizing error accumulation common in SfM methods when images of the entire surrounding area are used.

#### *MMM*: Multi-Masked Matching

In MMM, our proposed 3D point cloud generation method, a cloud is generated by stereo matching using images captured from closely positioned cameras locally (Multi-matching). This is followed by iterative point removal using mask images derived from the surrounding viewpoint images utilized during point cloud generation (Masked matching).

Figure 1 shows a representative example of the process. In this example, stereo matching is performed using two images from adjacent camera positions (indicated by the pink squares in Fig. 1A) to generate a 3D point cloud, referred to as the *baseCloud*. This process is executed sequentially using the Metashape API functions buildDepthMaps () and buildDenseCloud ().

**Figure 1.**
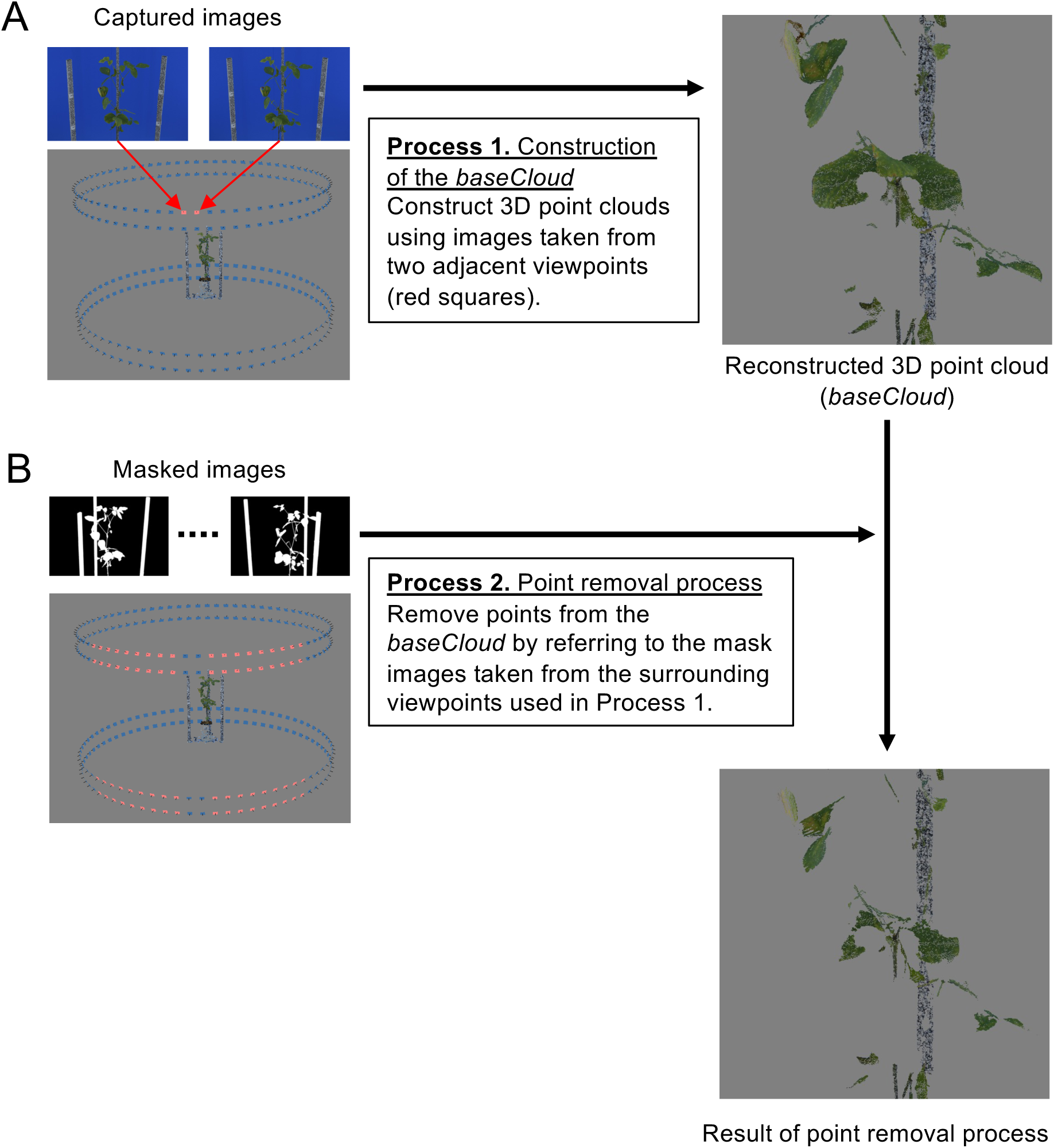
Algorithm of MMM (Multi-Masked Matching). A) Point clouds are generated locally using images from two adjacent viewpoints. B) Noise points are removed using mask images derived from the surrounding viewpoint images. Processes 1 and 2 are repeated for each viewpoint, and the resulting point clouds are integrated.

Next, mask images from other viewpoint images are used to remove noise points of the *baseCloud* (the 72 pink squares in Fig. 1B), taken from viewpoints within a 45-degree range to the left and right, facilitating the point removal process from the *baseCloud*. The mask image is a binary representation where the area of the plant region to be measured is set to 1, while other areas are designated as 0. The execution of this process involves utilizing the Metashape API functions selectMaskedPoints () and removeSelectedPoints () in that order. For every image captured, this process is carried out for all combinations of adjacent images to obtain the point cloud and remove noise points. The resulting point clouds are amalgamated to construct a 3D point cloud of the target plant.

#### *CLCFM*: Closed-Loop Coarse-to-Fine Method for SfM

We propose the CLCFM to improve the stability and accuracy of camera position estimation for images taken from all around the target. Instead of performing SfM simultaneously with all images captured around the target, we utilize a small subset of the images for the initial SfM process. We then iterate the process by incorporating subsets of the remaining images taken around the target. This iterative process employs a small number of all-around images initially, then incorporates all images to perform SfM to estimate the camera positions of all captured images. This method of performing SfM sequentially rather than all at once improves the accuracy of the camera position estimation.

The principle of the method is illustrated through an example where a single camera captures images of an object rotated in 5-degree increments (Fig. 2A). First, the images are categorized into groups, with the 72 images around the object being divided into six groups in this instance. Next, the order in which the SfM process is executed (GroupOrder) is determined. The GroupOrder is selected so as to minimize the bias in the estimated camera positions. For example, if Group 1 is the first group subjected to the SfM process, then Group 4, which includes the images halfway between the adjacent images of Group 1, is selected as the second group to be processed, and Group 2, which is between Group 1 and Group 4, is selected as the third.

**Figure 2.**
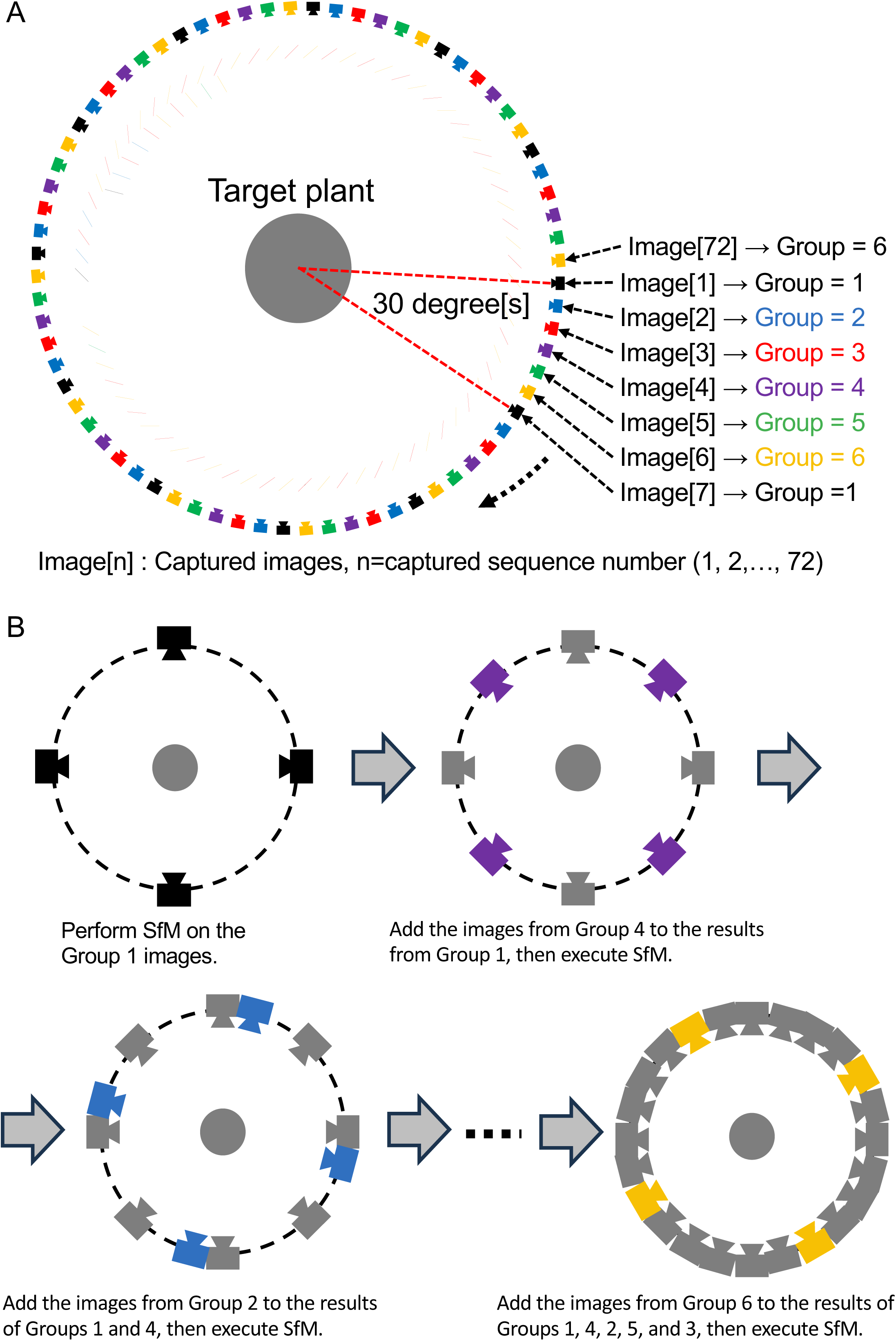
Algorithm of *CLCFM* (Closed-Loop Coarse-to-Fine Method). A) The sequentially captured images are divided into groups according to viewpoint. Six groups consist of 72 images taken at 5-degree intervals. B) Procedure of CFCFM. First, SfM is performed on group 1 images. Then, group 4 images are appended and SfM is performed. Similarly, groups 2, 5, 3, and 6 images are appended and SfM is performed.

Next, the procedure shown in Fig. 2B is followed to perform SfM sequentially for each group. The images in Group 1 are used to estimate the first camera position. The main process for camera position estimation in Fig. 2B, AlignProcess (), is executed for ImgList [] in the order of the Metashape API functions matchPhotos (), alignCameras (), and optimizeCameras (). The results of the first camera position estimation are then used as initial values, and a second SfM is performed by adding the Group 4 images to the Group 1 images. Similarly, images are processed in the order specified by GroupOrder, with SfM being sequentially executed using the camera position estimates obtained from the previous step as initial values. The final camera position estimates are derived by processing all groups. When images are captured using multiple cameras arranged vertically, those taken from the same vertical angle are grouped together for processing.

### 2.3. 3D point cloud reconstruction pipeline program

We developed a 3D point cloud reconstruction pipeline that implements our proposed method. This pipeline consists of four subprograms executed sequentially, starting with the captured image as the initial input to reconstruct the 3D point cloud (Fig. 3).

**Figure 3.**
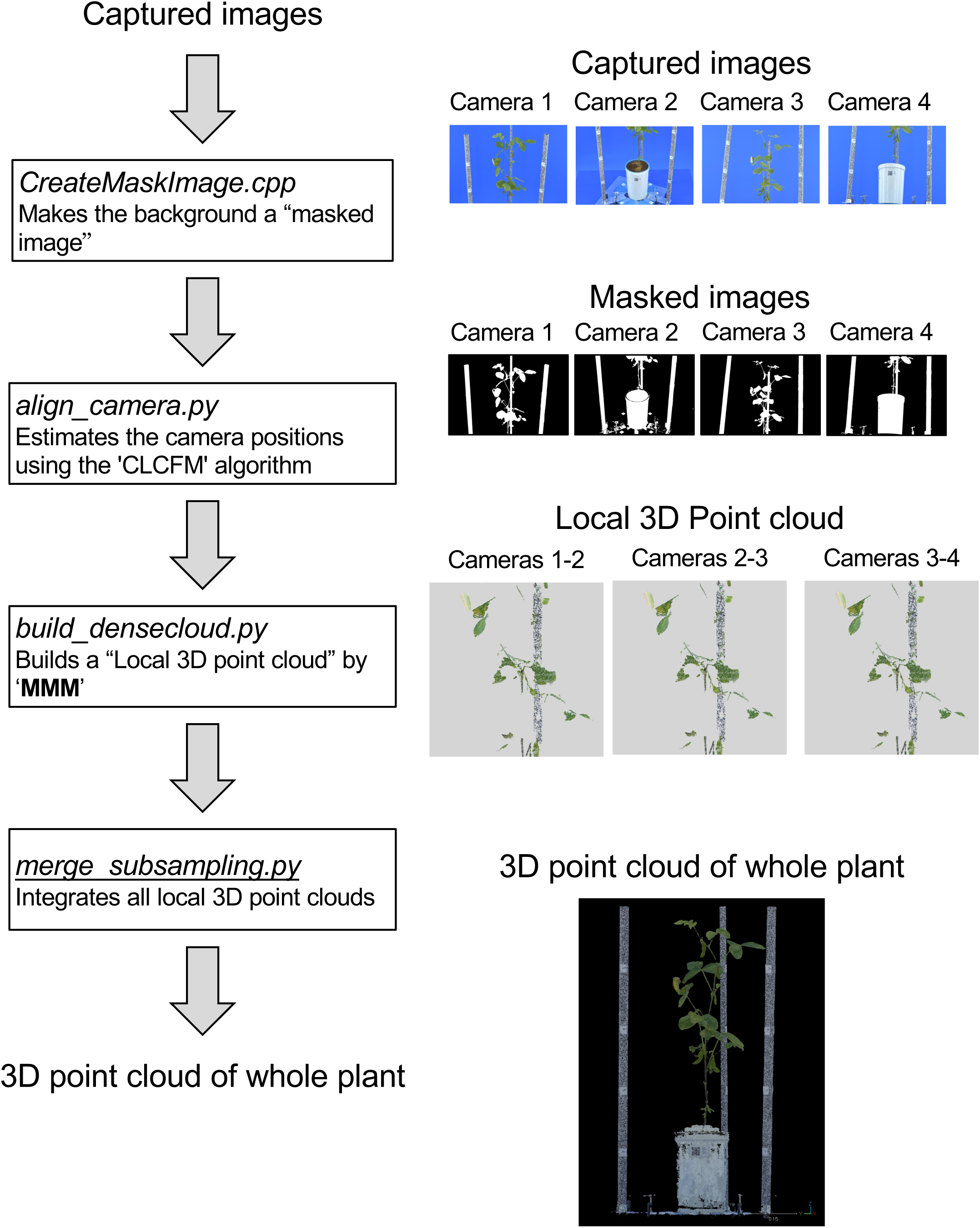
3D point cloud reconstruction pipeline program. ”CreateMaskImage.cpp” makes mask images. “align_camera.py” processes CFCFM. “build_densecloud.py” and “merge_subsampling.py” are MMM processes. These programs are executed sequentially with the captured image as the initial input to reconstruct the 3D point clouds.

1) CreateMask.cpp

This is a program for creating mask images. The program generates binary mask images where the plant areas vital for reconstructing the 3D point cloud and the relevant object regions are shown in white, while the remaining areas are black. Mask images are produced for all captured images, utilized for mask processing in CLCFM and noise elimination in MMM.

This program was developed using Visual C++ (Visual studio 2019) and OpenCV version 3.4.14.

2) align_camera.py

This script performs the camera position estimation process of the CLCFM algorithm, providing the camera position estimation information used in MMM.

This program was developed using Python 3.8, Metashape Professional Edition 1.8.5, and the Python 3 Module (Agisoft, https://www.agisoft.com).

3) build_densecloud.py

This script executes the MMM algorithm to construct a 3D point cloud and exports multiple 3D point cloud data files in PLY format. The script was developed using Python 3.8, Metashape Professional Edition 1.8.5, and the Python 3 Module. The results presented in this paper were obtained using the following execution parameters for the processing functions within the Python 3 Module.

matchPhotos(): The images were processed at 100% scale with accuracy set to ‘high’.
buildDepthMaps(): The images were processed at 1/4 scale (1500x1000 pixels) with quality set to ‘medium’.

4) merge_subsampling.py

This program integrates the diverse 3D point cloud data output by ‘build_densecloud.py’ and performs subsampling processing at 0.5 mm intervals, resulting in the output of the 3D point cloud of the target object.

This program was developed using Python 3.7 and open3D 0.15.

### 2.4. Imaging

To obtain the best images for the proposed CLCFM3 method, we built an imaging studio consisting of a rotating stage with plants, a camera, LED lighting, and background panels (Fig. 4A). The background panels, equipment, and cords were colored blue to facilitate the creation of mask images.

**Figure 4.**
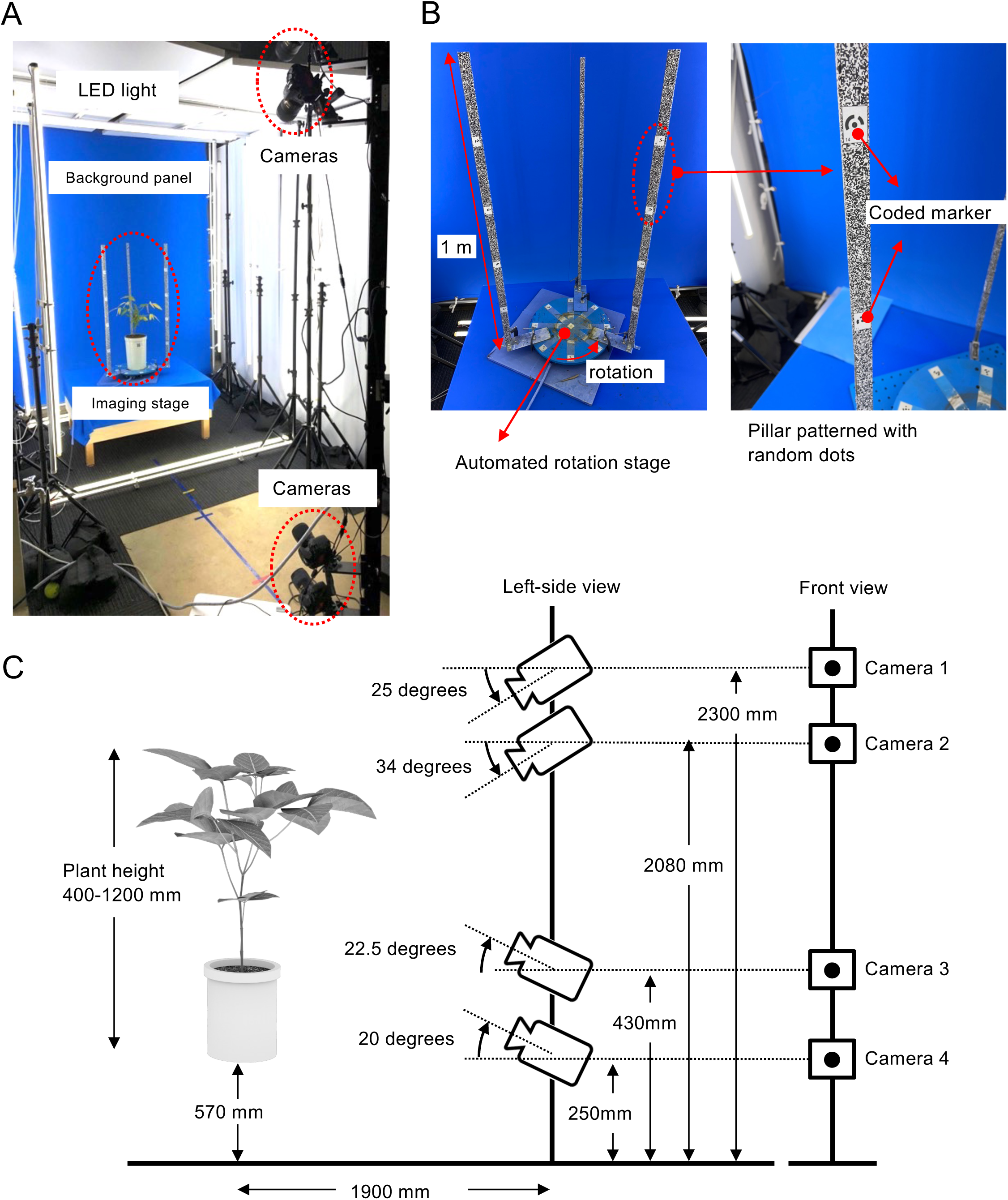
Imaging studio for 3D point cloud reconstruction by CLCFM3. A) The imaging studio consists of an imaging stage with a rotating platform on which plants are placed, four cameras positioned in front, LED lights surrounding the plant, and a blue background panel to facilitate mask image generation. B) The imaging stage has three pillars on which random dot patterns and coded targets for scale are attached. C) Camera angles and distance from the object are determined for soybeans ranging in height from 400 to 1200 mm.

The imaging stage was devised to allow the rotation of the plant in front of the camera, capturing a comprehensive image of the entire plant (Fig. 4B). SGSP-160YAW (Sigma Koki, Tokyo) and GIP-101 (Sigma Koki) were connected to a personal computer using a serial USB converter (COM-1(USB)H; Contec, Tokyo). The rotations were controlled by a program running on the personal computer.

To hold the cultivation pot, an acrylic disc (35 cm in diameter) was made and attached to the SGSP-160YAW. Small pieces of paper printed with coded targets were attached to the acrylic disc, and additional coded targets were attached to the three pillars fixed outside the perimeter of the disc. The coded targets were used as reference points for integrating the point cloud using MMM and to define the scale that gives the actual dimensions [6]. The coded targets were attached to the following positions: the center of the disc; eight positions at 45-degree intervals around the circumference, 15 cm from the center; and positions separated by 20 cm apart vertically on the pillars. Each of the three pillars consisted of an aluminum frame of 2.5 cm width Í 1.5 cm thickness Í1 m length, to which was attached a strip of paper printed with a random dot image. These random dots were used to ensure that the feature points obtained in the CLCFM and MMM processing steps were stable [6]. The pillars were attached to the side of the disc at uniform angles (Fig. 4B, left).

Images were captured using four digital SLR cameras (D5500; Nikon Corporation) and lenses (AF-S Micro NIKKOR 60mm f/2.8G ED, Nikon Corporation). Figure 4C shows the camera arrangement used to capture images of the plants used as targets with heights ranging from 400 to 1200 mm). The cameras were connected to a personal computer via USB cables. The shooting conditions were: shutter speed 1/15 sec, ISO sensitivity 100, aperture F/16. The acquired images was saved as 6000x4000 pixel JPG files.

## 3. Results

### 3.1. ‘*baseCloud*’ construction process in MMM

Figure 5 shows the results of an experiment to verify the optimal viewing angle for generating *baseCloud* point clouds of the target plant. Figures 5a and 5b display portions of the *baseCloud* reconstructed using only two adjacent images and four adjacent images, respectively, from different camera positions. These images were captured by rotating the table holding the object in 5-degree increments, resulting in viewing angles of 5 degrees for the *baseCloud* in Fig. 5a and 15 degrees for that in Fig. 5b. For this object, no significant difference was observed between the *baseClouds* in Figs. 5a and 5b. Nonetheless, while reconstructing a separate sample with Metashape 1.6, we confirmed that configuring the viewing angle to 5 degrees helped reduce the loss of point clouds (Fig. S3). The reconstruction time with MMM diminishes with a reduction in the number of images employed. Given these findings, we chose to generate the *baseCloud* using two adjacent images.

**Figure 5.**
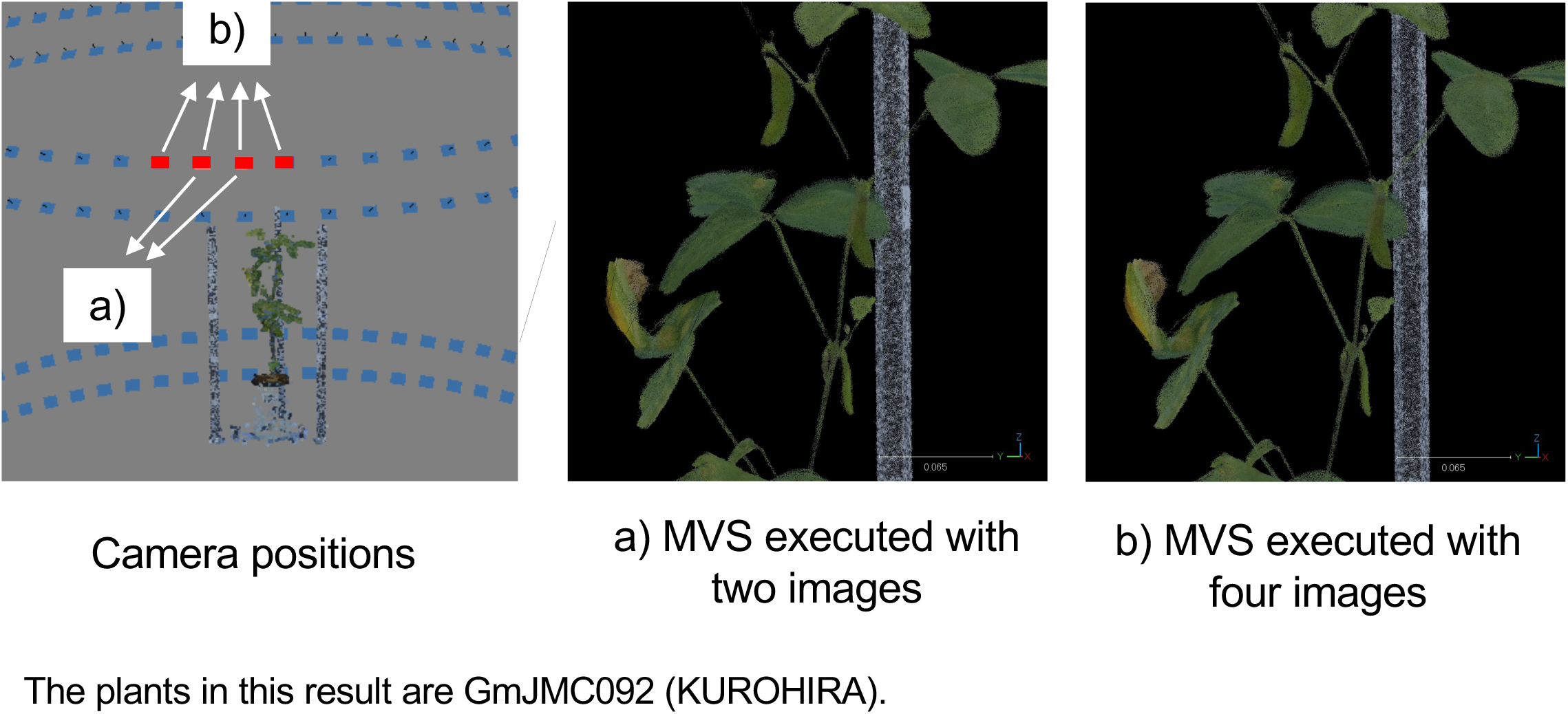
Results of ‘baseCloud’ reconstruction with different numbers of images. a) Two adjacent images. b) Four adjacent images. Processing time increases with the number of images, while there is no significant difference between a) and b).

### 3.2. Viewing angle of the mask image used in the MMM point removal process

We next investigated the optimal viewing angle for the mask image used in the point removal procedure of the *baseCloud* performed by MMM. To ascertain the optimal angle, we performed the point removal process of *baseCloud* using MMM under four different viewing angles of the mask image, similar to the method illustratedin Fig. 6. Figure 6 illustrates the outcomes when employing the mask image at viewing angles of a) 50 degrees, b) 90 degrees, c) 180 degrees, and d) 360 degrees. The results indicated that as the viewing angle of the surrounding mask increased, more points were eliminated; however, this increase also led to the deletion of points in the plant body that should have been retained. We therefore decided to set a 90-degree viewing angle of the mask image used in point removal.

**Figure 6.**
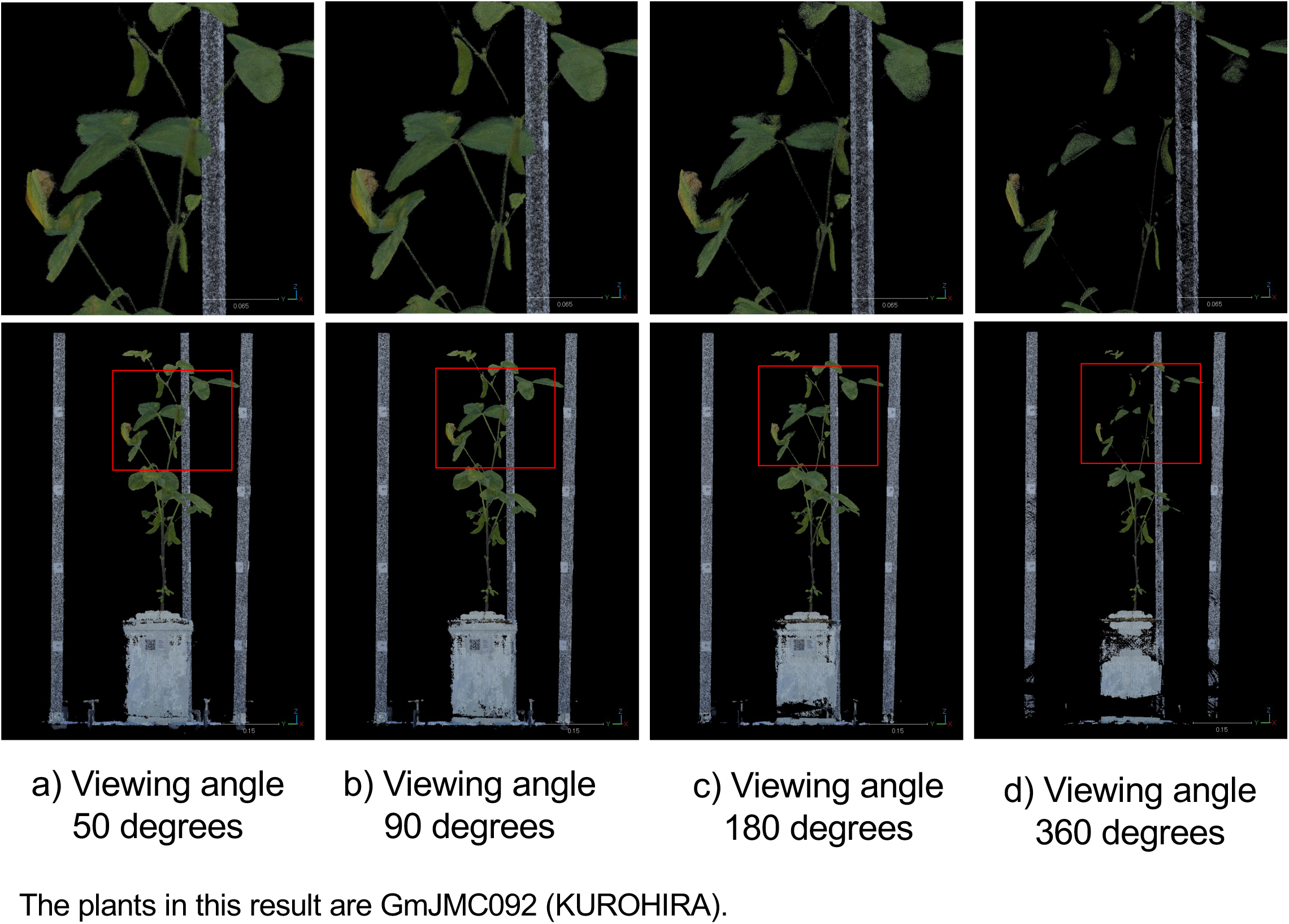
Results for different viewing angles of the mask image used in the point removal process. a) 50 degrees, b) 90 degrees, c) 180 degrees, d) 360 degrees. More points are deleted as the viewing angle increases, while points consisting of leaves and stems are also deleted.

### 3.3. 3D point cloud reconstruction of soybean plants

Using the pipeline developed and the processing conditions established in the previous sections, we successfully reconstructed 3D point clouds for four soybean varieties (Fig. 7a, Fig. S1). The results demonstrated the proposed method’s ability to reconstruct 3D point clouds of varying shapes, sizes, and structural complexities. For comparison, the identical image sets underwent standard Metashape functions to generate 3D point clouds. This comparison confirmed that the proposed method effectively mitigated the appearance of blue dots as noise in the outlines of leaves and stems, while also resolving the problem of point clouds failing to form in certain thin-stem regions (Fig. 7b, Fig. S1, Fig. S2).

**Figure 7.**
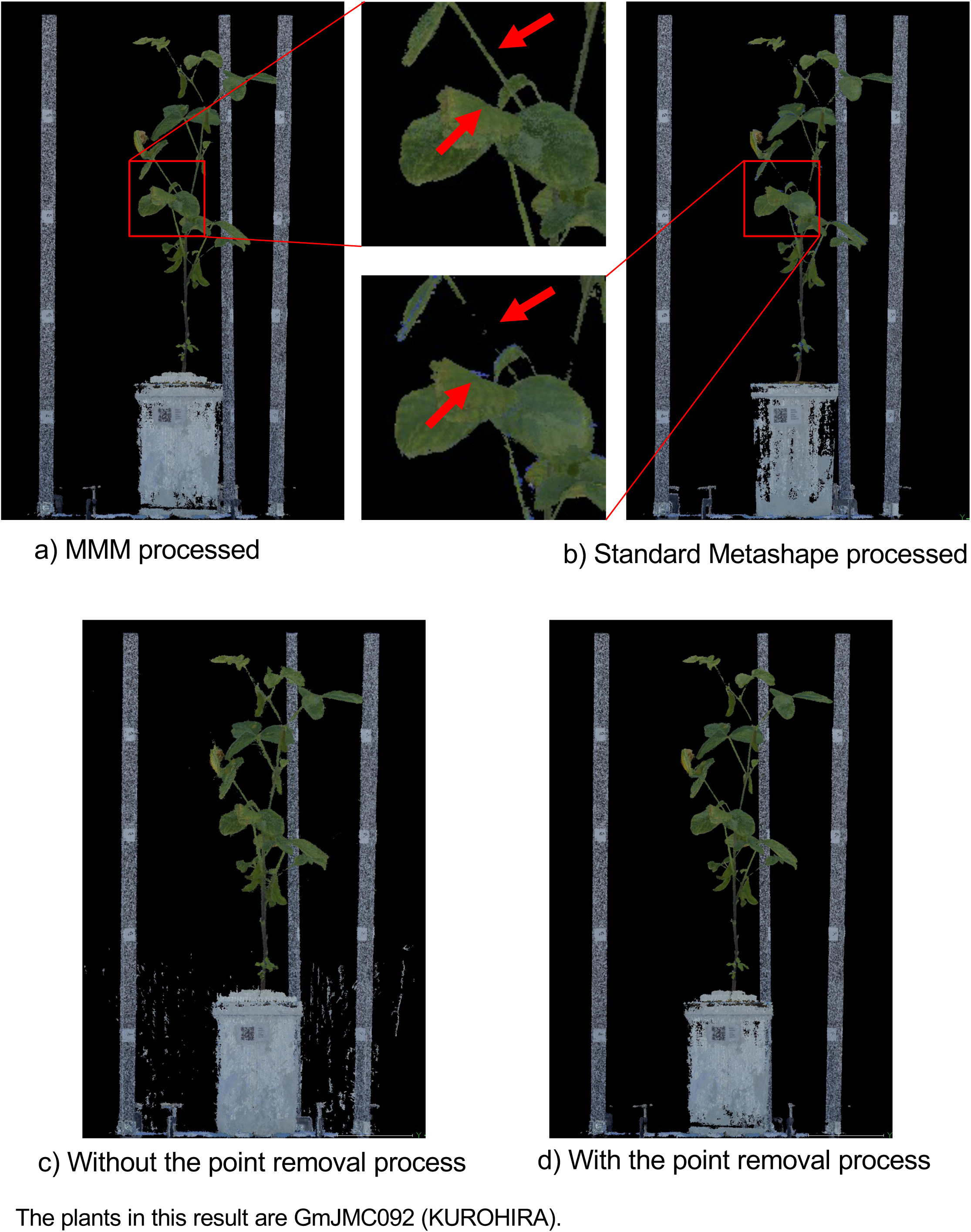
Differences in results between point removal processes. a) and d) Removed by MMM. b) Standard Metashape process. c) Removed by MMM without the point removal process.

Figure 7c displays the outcome when the point removal process was omitted after the *baseCloud* was generated with MMM, while Fig. 7d showcases the result of the point removal process after the *baseCloud* was generated with MMM. In both cases, point cloud reconstruction and point removal processing were performed after SfM was executed using CLCFM. Without point removal processing, noise points emerged in the background and other parts of the captured image (Fig. 7c), while point removal processing resulted in the removal of noise points (Fig. 7d).

### 3.4. Effects of the point removal processes in MMM

We compared the differences in the reconstruction of 3D point clouds by MMM with and without point removal processing. The *Base clouds* were generated using the MMM point cloud reconstruction approach, leading to the production of 3D point clouds after applying point removal processing (Fig. 8a) and without such processing (Fig. 8b). These 3D point clouds were then overlaid and compared within a virtual space. To facilitate this comparison, only the plant body was extracted, while the support, pot, and rotating stage were manually removed. The distance between corresponding points in the aligned 3D point clouds was calculated. Points with distances of 3 mm or less were deleted, yielding a point cloud that represented the differences between the two 3D reconstructions (Fig. 8c). This process was repeated for distance intervals of 0–1 mm, 1–2 mm, 2–3 mm, and 3–22 mm, with the number of points constituting each point cloud recorded (Fig. 8d). The analysis revealed that in the point cloud post-point removal process, 78,637 points exhibited a distance variance of 3 mm or more, constituting 4.3% of the aggregate points within the point cloud (Fig. 8d).

**Figure 8.**
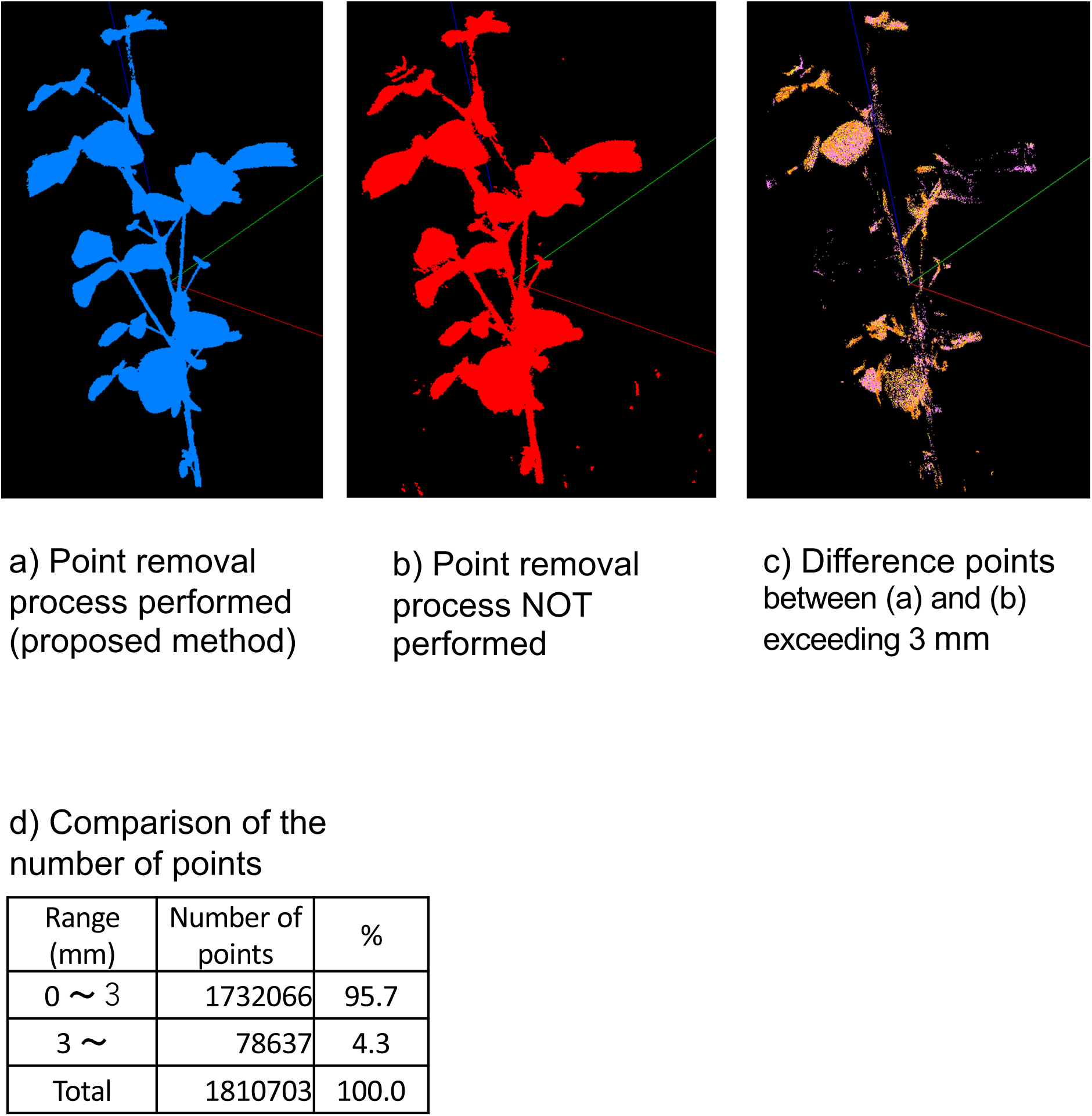
Results comparing point cloud reconstruction with and without the noise removal process by MMM. a) Point clouds with noise removal process by MMM. b) Point clouds without noise removal process by MMM. c) Point clouds where the distance between a) and b) is 1-3 mm. d) Number of point clouds categorized by distance ranges.

### 3.5. Validation of position accuracy in the reconstructed point cloud

We assessed the position estimation errors for the 3D point cloud acquired through the CLCFM3 pipeline using the evaluation model shown in Figure 9. The evaluation model was arranged with 13 pairs of measurement points (Fig. 9 (1)-(13)), consisting of coded target pairs. The distance between these targets was measured manually with a caliper and compared with the value measured using the 3D point cloud generated by CLCFM3. The actual values were measured 10 times each, and the average was used. The coded targets attached to the pillars were measured using a ruler to derive the average value (Fig. 9 (14)). The reading errors for manual measurement were set at 0.02 mm for the caliper and 1 mm for the ruler. The measurement using CLCFM3 was acquired by obtaining the distances between the targets in the 3D point cloud reconstructed through the pipeline. This procedure was repeated 10 times, and the average value was derived.

**Figure 9.**
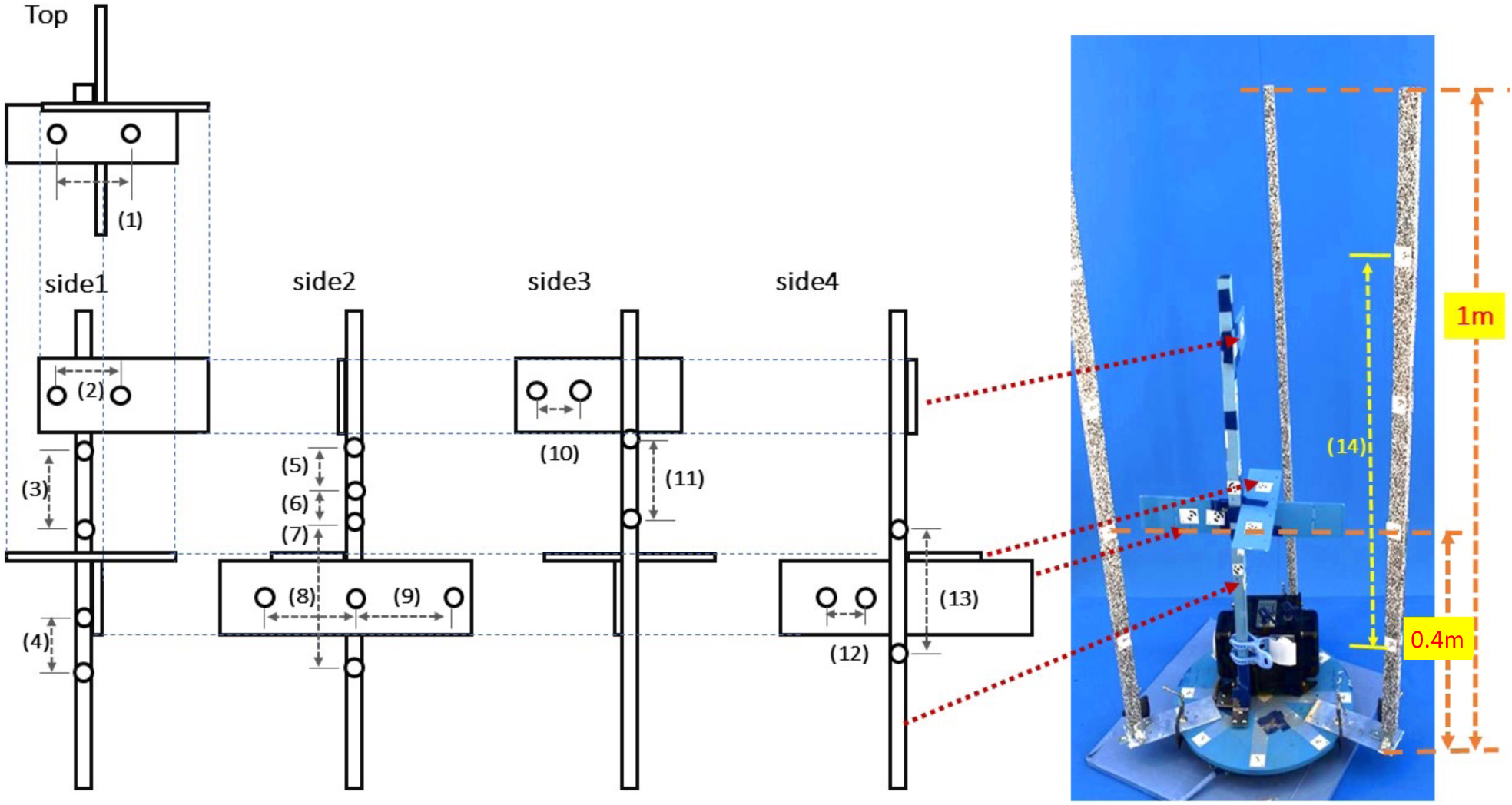
Model for evaluating accuracy and measurement positions. CLCFM3 is processed 10 times individually, and the distances of measurement positions (1)-(14) are measured respectively.

Table 1 shows the results. For measurement points (1) to (13), the maximum difference between the manually measured values (actual values) and the values obtained from the 3D point cloud was 0.280 mm, with a maximum standard deviation of 0.0000346 mm for the 3D measurements.For measurement point (14), the difference between the average manually measured value (actual value) and the average value obtained of the measured values was -0.97 mm, and the standard deviation of the measured values was 0.00012 mm. For measurement point (14), the difference between the average manually measured value (actual value) and the average value obtained from the 3D point cloud was -0.97 mm, with a standard deviation of 0.00012 mm for the 3D measurements.

**Table 1.** Evaluation of measurement accuracy by the proposed pipeline. Caliper values are obtained by measuring the length 10 times using a caliper for (1)-(13) and a ruler for (14), and then calculating the average. CFCFM3 values are obtained using the point cloud data obtained using CLCFM3. “Difference” is the difference in distance between the caliper and CFCFM3.

| Measurement position |  | Caliper<br>(mm) | CFCFM3 |  | Difference<br>(mm) |
| --- | --- | --- | --- | --- | --- |
| Number | Direction |  | Avg(mm) | SD |  |
| (1) | width | 124.24 | 124.352 | 0.0000314 | -0.112 |
| (2) | width | 84.51 | 84.651 | 0.0000346 | -0.141 |
| (3) | height | 120.05 | 120.104 | 0.0000271 | -0.054 |
| (4) | height | 69.15 | 69.160 | 0.0000068 | -0.010 |
| (5) | height | 63.11 | 63.151 | 0.0000115 | -0.041 |
| (6) | height | 42.45 | 42.356 | 0.0000110 | 0.094 |
| (7) | height | 156.36 | 156.357 | 0.0000260 | 0.003 |
| (8) | width | 131.41 | 131.130 | 0.0000225 | 0.280 |
| (9) | width | 133.6 | 133.509 | 0.0000176 | 0.091 |
| (10) | width | 44.27 | 44.356 | 0.0000099 | -0.086 |
| (11) | height | 101.05 | 101.183 | 0.0000201 | -0.133 |
| (12) | width | 47.6 | 47.793 | 0.0000067 | -0.193 |
| (13) | height | 144.38 | 144.579 | 0.0000191 | -0.199 |
| (14) | height | 593 | 593.970 | 0.0001225 | -0.970 |

## 4. Discussion

### 4.1. Generating local 3D points and removing noise with mask images

We proposed an approach to reduce the loss due to occlusion by constructing 3D point clouds using images with narrow viewing angles. However, this process generates many flying pixels due to errors in stereo matching when searching for corresponding points along the outline. We proposed to solve these issues by eliminating points using a mask image of the peripheral field of view from the images employed in point cloud reconstruction, reducing occlusion-related losses and achieving point cloud reconstruction with fewer noise points.

Figure 10 shows a schematic of the image-matching process at a plant boundary.

**Figure 10.**
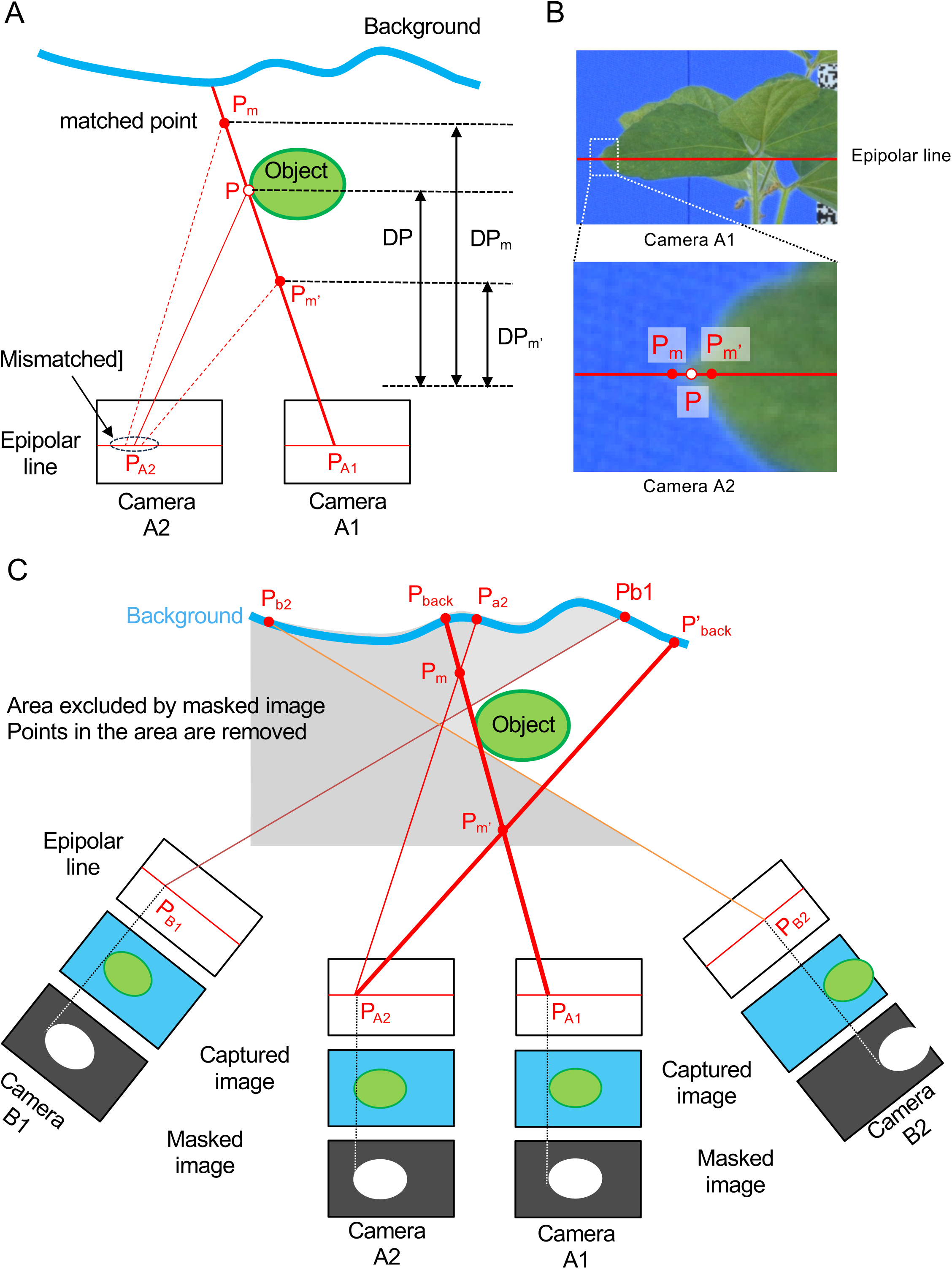
Image-matching process at the boundary. A) Process of mismatching. B) Difference between correct point and mismatched point. C) Point removal process using masked images derived from surrounding viewpoint images (Masked Matching).

To estimate the 3D position of point P on the object’s contour line, we extract the coordinates of point P_A1_ and point P_A2_, representing point P’s location in the images taken by cameras A1 and A2, respectively. Using these coordinates and the camera position information for cameras A1 and A2 acquired through SfM, the depth distance DP between point P and the cameras is calculated. Based on the outcome, the 3D position of point P is ascertained, leading to the creation of a point for the 3D point cloud (Fig. 10A).

Here, we focus on point P in the outline of the plant body. Image matching is performed to find position P_A2_ on the image taken by camera A2 relative to position P_A1_ on the image taken by camera A1.

The DP value fluctuates according to the matching results. If the position of point P found by camera A1 is correctly matched with point P in the image from camera A2, DP is assigned is accurate value. Consequently, point P on the plant body will be generated in 3D space (Fig. 10A). However, if point P is incorrectly matched and the matching result differs from the accurate point P (Fig. 10B), the DP value deviates, leading to the acquisition of DP_m_ and DP_m’_ values (Fig. 10A). Based on these values, points are generated at positions Pm and Pm’, which differ from the accurate position of point P. These points, P_m_ and P_m’_, are noise points that do not exist on the plant body. The causes of inaccurate image matching include the resolution of the imaging camera’s image sensor and the matching process with points exhibiting textures akin to leaves and stems close to point P. To address the issue of noise points caused by mismatching, our proposed method does not aim to rectify the mismatches but instead identifies numerous candidate points, incorporating a post-processing step for point removal. This approach reduces the number of noise points.

Figure 10C shows a schematic of the point removal process for MMM. Cameras A1, A2, B1, and B2 represent the camera positions. The A1 and A2 images are used to generate the point cloud (*baseCloud*), and the mask images generated from the images captured by all four cameras are used for the point removal process (Fig. 10C). The mismatching at points on the contour line of the object occurs in the vicinity of the intersection lines P_A1_P_back_ and P_A2_P’_back_ of Cameras A1 and A2. Using the A1 and A2 mask images, the point removal process can eliminate mismatched points beyond the object’s boundaries. However, none of the mismatched points generated on the P_A1_P_back_ and P_A2_P’_back_ boundary lines can be removed. The mismatched points P_m_ and P_m’_ on the boundary line shown in Fig. 10C can be eliminated by excluding areas outside the object at the intersection lines P_B1_P_b1_ and P_B2_P_b2_ of Cameras B1 and B2 in the mask image. This approach also enables the removal of mismatched points along the line between P_A1_P_back_ and P_A2_P’_back_.

The proposed method uses surrounding mask images to detect flying pixel noise points. Therefore, the accuracy of noise point detection depends on the results of the mask image region extraction, making the image processing of the region extraction an important factor. The developed pipeline employs a method that uses color differences for region extraction [29, 30]. Numerous image analysis techniques have been proposed for area extraction, enabling the generation of tailored mask images capable of eliminating noise points adapted to various plants and shooting environments. This versatility allows MMM to be applied across a wide range of plants, facilitating the reconstruction of 3D point clouds with suppressed flying pixel noise.

The proposed method can remove noise points on the outside of the plant body using a point removal process. However, noise points located inside the plant body cannot be detected and removed. Developing a method for removing these internal noise points stands as a future issue.

### 4.2. Enhancing camera position accuracy for all-around images

In this study, we used all-around images taken by rotating the object at 5-degree intervals for 3D point cloud reconstruction. In estimating camera position using SfM processing on omnidirectional images with narrow angles of view, one challenge is that the estimation process is likely to fail due to accumulated errors (Fig. 11A).

**Figure 11.**
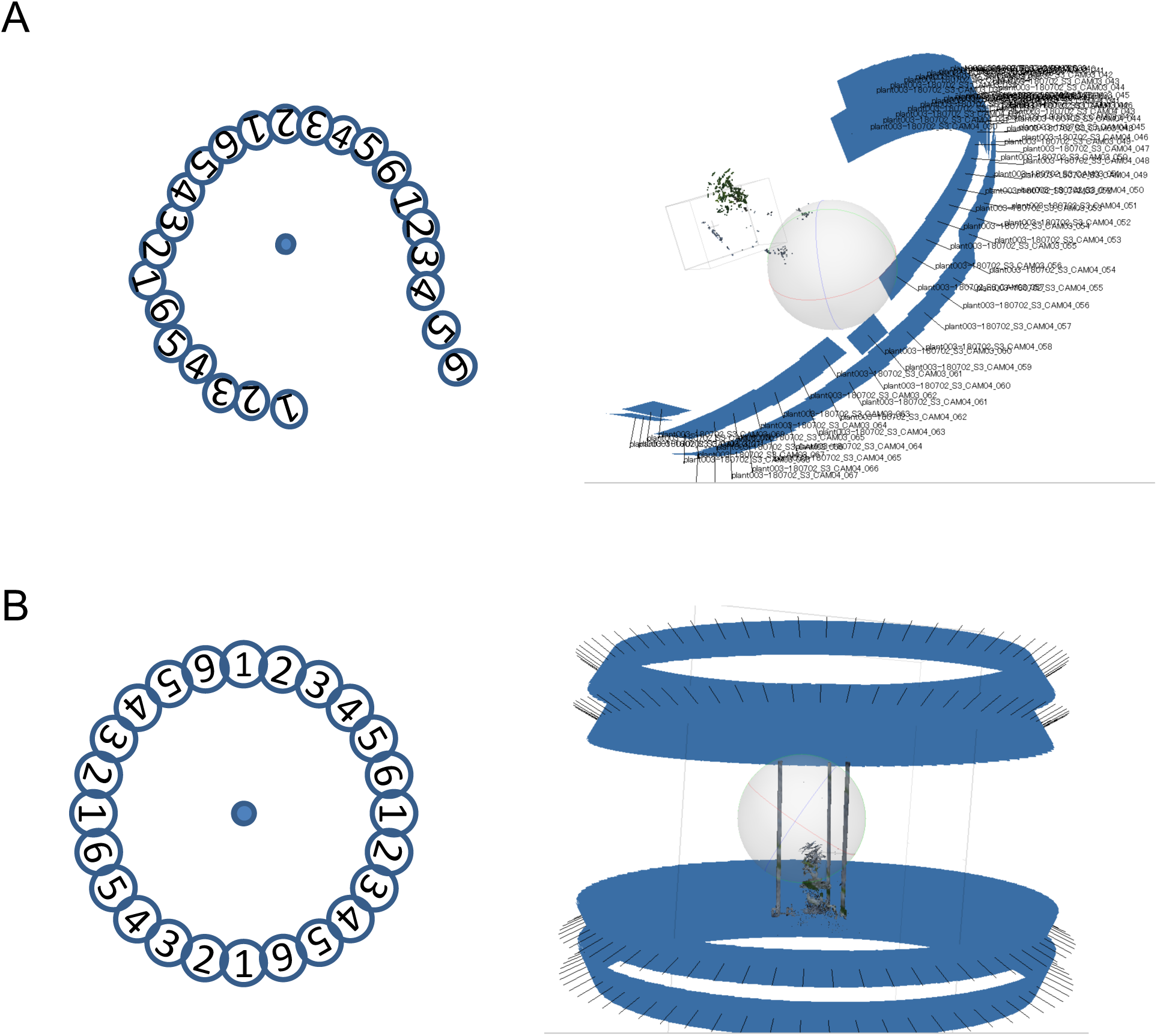
Camera position estimation failure. There is a risk of camera position estimation failure due to the accumulation of camera position estimation errors for all-around images taken at short intervals in SfM. A) Without CFCFM and pillar. B) With CFCFM and pillar.

CLCFM estimates camera position while gradually adding panoramic images. First, camera position is estimated using only images that surround the object but have an open viewpoint, and the results are used as initial values. Next, we add a few more images and run the estimation process again, updating the results. We reiterate this process until we have estimated the camera positions for all the images in the full surround. This processing method is expected to reduce the failure of position estimation due to error accumulation by suppressing the propagation of estimation errors that occur in the estimation process (Fig. 11B).

However, there is a problem with implementing this algorithm. If the cameras are placed far apart in the initial stage of processing, there are few common feature points between the images, making it difficult to estimate camera position in the initial processing. We solved this problem by affixing random patterns to the pillars when taking pictures (Fig. 4B). Securing numerous image feature points within the area featuring the random pattern allowed for a bundle adjustment with self-calibration [31]. This made it possible to stably obtain the camera’s internal and external parameters. Finally, since it is not necessary to perform internal and external calibration in advance, a key advantage is the potential to attain high measurement accuracy without engaging in device setup or maintenance [6].

## 5. Conclusion

This paper introduced the CLCFM3 algorithm for the 3D reconstruction of all-around plant images. Given that plants possess a distinctive morphology characterized by overlapping thin parts such as leaves and stems, challenges arise when performing 3D point cloud reconstruction via photogrammetry. These challenges include the inability to acquire points in the 3D point cloud due to occlusion in the overlapping parts and the generation of points where the object does not exist due to image matching errors along the object’s outline.

To solve these problems, we proposed the MMM approach. MMM generates a point cloud using images taken from a viewpoint close to the object to avoid occlusion (multi-matching) and then repeatedly executes point cloud removal processing using surrounding mask images to remove the noise points (mask matching) (Fig. 1).

To execute MMM, a highly accurate camera position estimation must first be obtained by SfM processing. To meet this requirement, we proposed CLCFM, which estimates camera positions with high accuracy for images encompassing the entire surroundings of the target. However, when SfM is conducted using such images, estimation errors accumulate. CLCFM selects images for SfM processing and iterates the SfM procedureto suppress this accumulation of errors (Fig. 2). MMM and CLCFM are based on the assumptions that the target is rotated and photographed from all sides and that the order of the images is known. Using this information, we were able to implement the proposed method by reconstructing local point clouds and selecting surrounding mask images for point removal processing.

We built an imaging studio (Fig. 4) and an image analysis pipeline (Fig. 3) to implement the proposed method and reconstructed 3D point cloudsfor soybean plants (Fig. 7). We validated the feasibility of reconstructing a high-density 3D point cloud of a plant body that minimizes the loss of thin stems, a characteristic shape of the plant body, as well as reducing noise from 3D points generated in areas devoid of plant structure (Fig. 8).

In this paper, Metashape Professional Edition 1.8 was used to execute SfM/MVS. We found that the different versions of Metashape yielded different results in MVS point cloud reconstruction (Fig. S3, Fig. S4). However, regardless of the version, the proposed method is effective for reconstructing 3D point clouds of plant bodies characterized by overlapping organ shapes such as leaves and stems. This effectiveness is based in the method’s ability to reduce noise by through point cloud reconstruction using local perspectives with MMM and point removal procedures involving mask images.

Although numerous methods for constructing 3D models of plants have been proposed, practical applications for measurement purposes remain limited except for rough assessments of community dynamics in the field. The proposed method enables the robust construction of highly accurate 3D models, a feature anticipated to advance the widespread use of 3D modeling for plant measurements in plant research.

## Supplementary Materials

Figure S1 Results of reconstruction of 3D point clouds of four soybean varieties using the proposed method.

Figure S2 Results of reconstruction of 3D point clouds for the image in Fig. S1 using Metashape

Figure S3 Results of running the analysis in Fig. 5 using Metashape 1.6.

Figure S4 Results of running the analysis in Fig. 6 using Metashape 1.6.

## Acknowledgments

**General:** We would like to thank Keiko Saegusa, Rumi Fujihira, Satomi Nakayama, Chie Kasashima, and Sayaka Kikuchi for their technical support at the Kazusa DNA Research Institute.We also thank Prof. Ryo Akashi, Dr. Hidenori Tanaka, Dr. Masatsugu Hashiguchi, and Dr. Takuyu Hashiguchi for providing soybean materials in this study.

**Funding:** This work was supported by a CREST Grant (No. JPMJCR16O1) from the Japan Science and Technology Agency and the Kazusa DNA Research Institute Foundation.

## Author contributions

HA, KN, IS, and TT contributed to the design of the study. HA developed the 3D reconstruction pipeline programs. HA and TT made imaging hardware. HA, KN, and KK conducted experiments and analyzed and interpreted data. HA, KN, IS, and TT wrote and revised the manuscript. KN and TT supervised the study.

## Data availability

The computer codes and scripts of MMM and CLCFM are deposited in a GitHub repository at https://github.com/tanasoft/MMM-CLCFM. Download links to image and 3D point cloud data that support the findings of this study are also provided at this GitHub repository.

---

**Supplemental Fig. S1.**
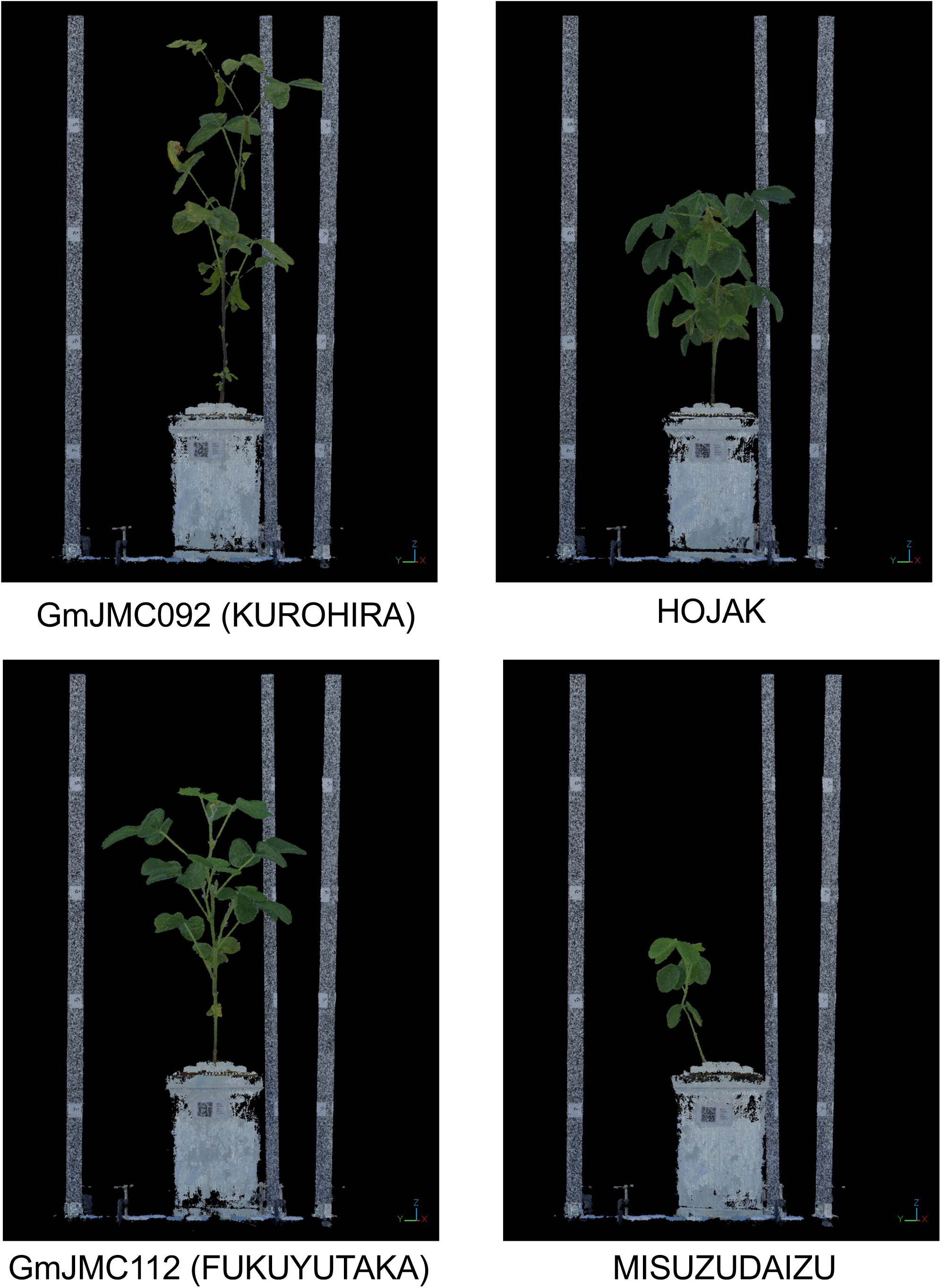
Results of reconstruction of 3D point clouds of four soybean varieties using the proposed method. The PLY data in this figure can be downloaded at https://github.com/tanasoft/MMM-CLCFM

**Supplemental Fig. S2.**
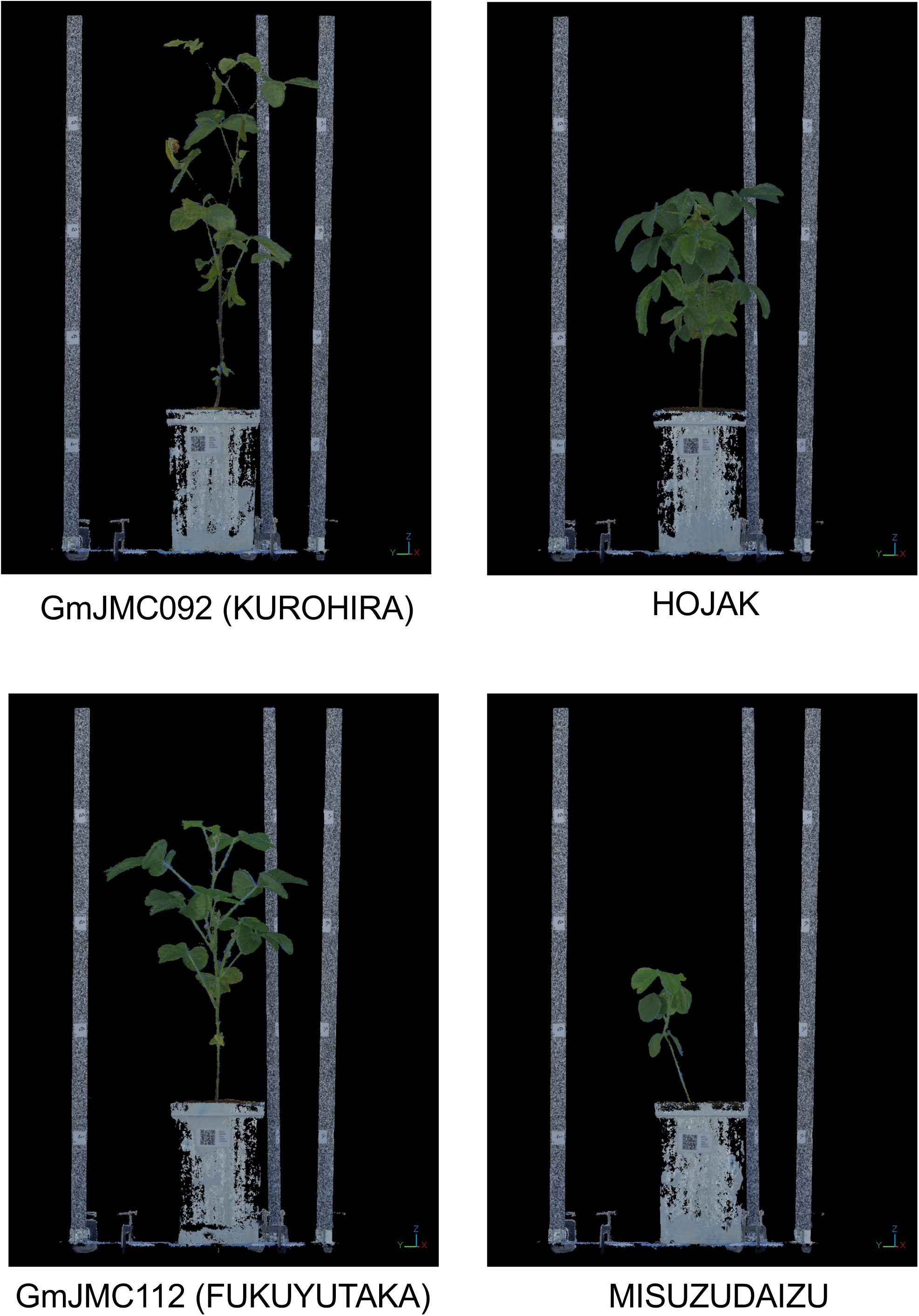
Results of reconstruction of 3D point clouds for the same image in Fig. S1 using Metashape.

**Supplemental Fig. S3.**
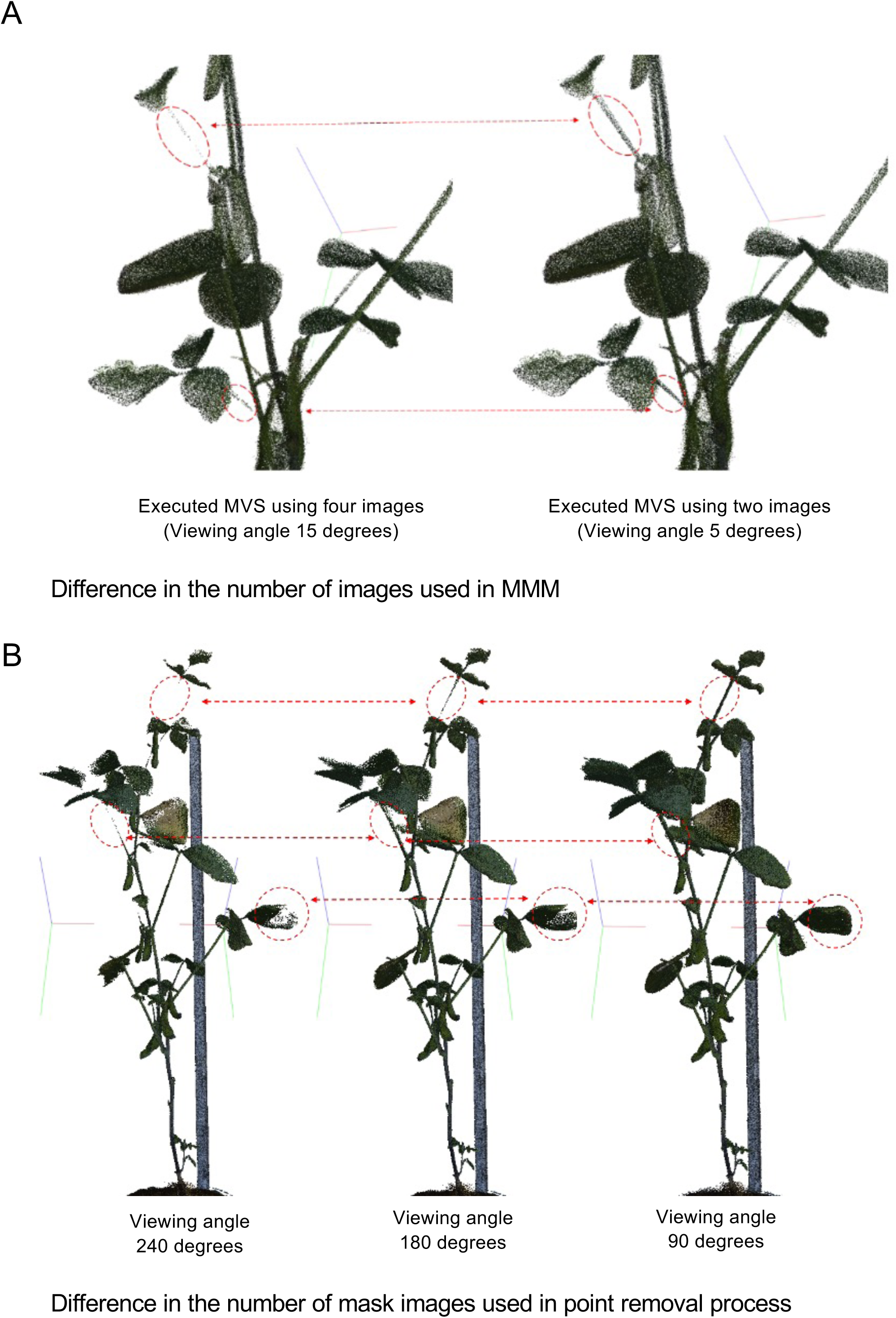
Results of running the analysis in Figs. 5 and 6 using Metashape 1.6.

**Supplemental Fig. S4.**
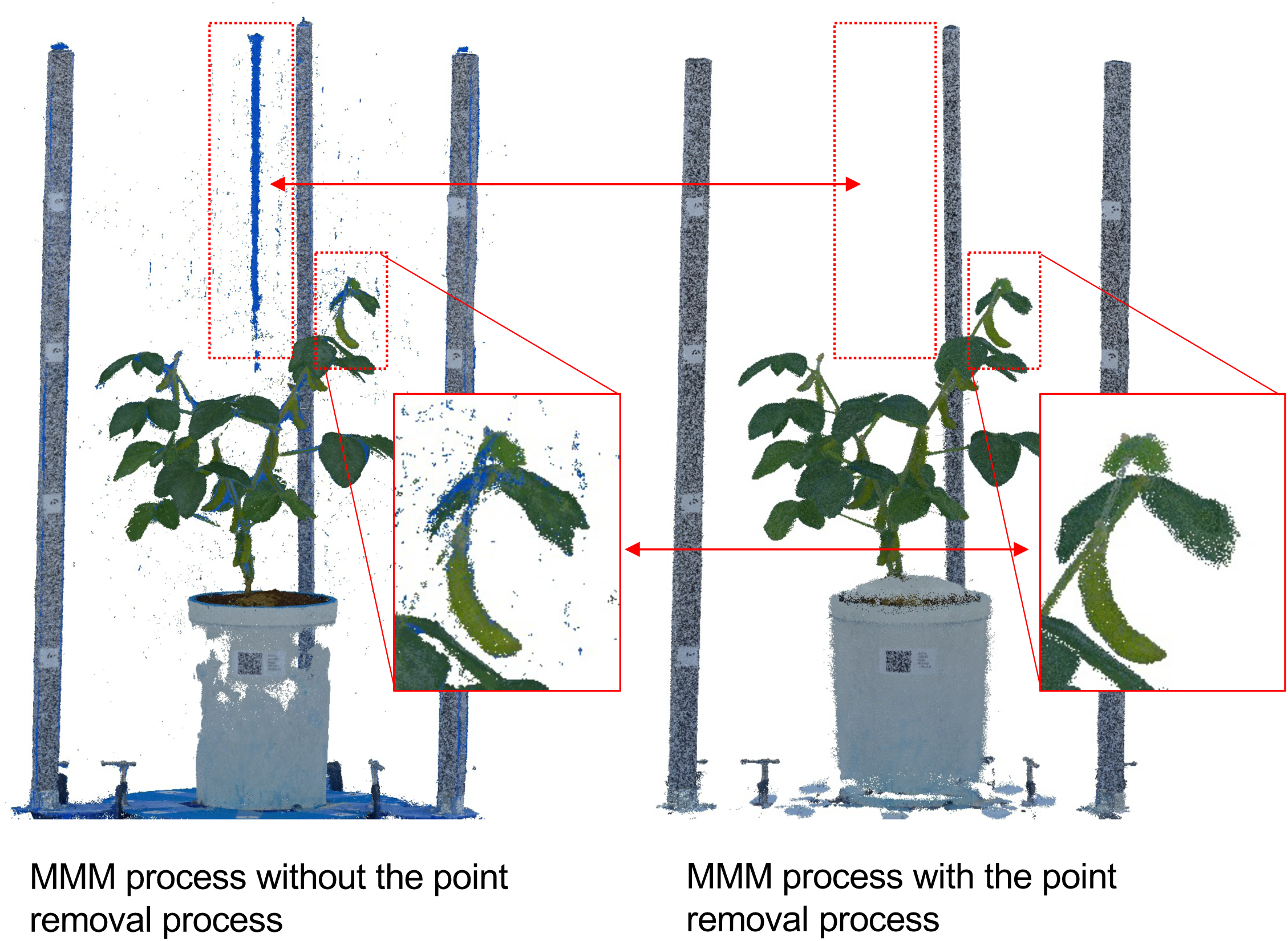
Results of running the analysis in Fig. 7 using Metashape 1.6.

